# Stimulating the medial prefrontal cortex disrupts inhibitory control over memory by modulating frontal and parietal brain regions

**DOI:** 10.1101/2024.01.30.577598

**Authors:** Ahsan Khan, Chun Hang Eden Ti, Kai Yuan, Maite Crespo Garcia, Michael C. Anderson, Raymond Kai-Yu Tong

**Author notes:** **Corresponding authors:** Raymond Kai-Yu Tong, Michael C. Anderson.

## Abstract

The act of recalling memories can paradoxically lead to the forgetting of other associated memories, a phenomenon known as retrieval-induced forgetting (RIF). This effect is thought to be mediated by inhibitory control mechanisms in the prefrontal cortex of the brain. Here we investigated whether stimulation of the medial prefrontal cortex (mPFC) with transcranial direct current stimulation modulates inhibitory control during memory retrieval in a RIF paradigm. In a randomized study, fifty participants received either real or sham stimulation, before performing retrieval practice on target memories. After retrieval practice, a final test was administered to measure the impact of stimulation on RIF. We found that stimulation selectively increased the retrieval accuracy of non-target memories and thus decreased RIF, suggesting a disruption of inhibitory control. Meanwhile, no change arose for the retrieval accuracy of target memories. The reduction in RIF was caused by a more pronounced beta desynchronization within the left dorsolateral prefrontal cortex (left-DLPFC), in an early time window (<500 msec) after the onset of the cue during retrieval practice. This, in turn, led to a stronger beta desynchronization within the parietal cortex in a later time window, an established marker for successful memory retrieval. Together, our results establish the causal involvement of the mPFC in actively suppressing competing memories and we demonstrate that while forgetting arises as a consequence of retrieving specific memories, these two processes are functionally independent. Finally, we demonstrate that beta desynchronization in the fronto-parietal brain regions indicates the disruption of inhibitory control.

## Introduction

The act of recalling a memory may not only facilitate the retrieval of intended information but can also induce forgetting of competing non-target memories, a phenomenon referred to as retrieval-induced forgetting (RIF) ((Anderson et al., 1994); see (Anderson, 2003; Anderson and Hulbert, 2020; Marsh and Anderson, 2022) for reviews). Despite extensive research efforts in the last two decades, the underlying psychological and neurobiological mechanisms of RIF remain only partially understood. This study investigates these mechanisms by applying stimulation to a key region within the episodic memory network, namely the medial prefrontal cortex (mPFC), and examining its impact on behavioural and electrophysiological indicators of RIF.

RIF arises in diverse materials, including words (Anderson et al., 1994), languages (Levy et al., 2007), and visual scenes (Shaw et al., 1995). Typically, the retrieval practice paradigm is used to study RIF, which consists of three phases (Anderson et al., 1994). In an initial study phase, participants learn category-exemplar pairs from several categories for a later memory test. In a subsequent retrieval practice phase, half of the items from a subset of the studied categories are retrieved repeatedly. A final test phase then occurs in which participants are tested on all of the studied items. Memory for the practiced items is enhanced compared to non-practiced items from non-practiced categories (i.e., control items), reflecting a memory facilitation effect (FAC). Critically, however, final recall performance for non-practiced items from practiced categories - items that would have directly competed with practiced items during retrieval practice trials is reduced compared to recall performance for control items. The tendency for retrieval practice to impair retention of competing items reflects the RIF phenomenon, a form of active forgetting.

Inhibitory control processes are thought to contribute to RIF (Anderson, 2003; Storm and Levy, 2012; Murayama et al., 2014; Anderson and Hulbert, 2020; Marsh and Anderson, 2022). According to the inhibitory model, two processes contribute to selectively retrieving a target trace in the face of competition from alternative memories during retrieval practice. First, when retrieval cues appear, a retrieval process automatically activates all traces in memory associated to those cues. Second, in response to this diverse activation, an inhibitory control mechanism is recruited to resolve interference from non-target traces by inhibiting those competitors, facilitating target retrieval. The two processes posited by the inhibitory control model necessarily interact. For example, for activation of competing memories to trigger inhibitory control, some mechanism needs to detect emerging competition that drives an increase in control. By this inhibitory model, RIF reflects the persisting aftereffects of inhibition on competing memories. An alternative explanation instead suggests that retrieving target memories strengthens those retrieved items, and this dominance causes them to block access to competitors on the final test (for a review of alternatives, see (Anderson et al., 1994; Anderson, 2003)). Thus, the inhibition and blocking views diverge on the relationship between FAC and RIF: whereas the non-inhibitory model directly ties the magnitude of RIF to the amount of FAC, the inhibitory model posits that these phenomena are functionally independent.

A large body of research on inhibitory control in attention involving the Stroop, Flanker and Simon tasks suggests that the anterior cingulate cortex (ACC) that sits in the mPFC detects the conflict and upregulate lateral prefrontal cortex (LPFC) activity to resolve it (e.g., (Hanslmayr et al., 2008; Kim et al., 2014; Crespo-García et al., 2022)). This same conflict detection mechanism is thought to detect competition during memory retrieval (e.g., (Kuhl et al., 2007; Crespo-García et al., 2022)) and to dynamically adjust as conflict is reduced. For example, using the RIF procedure, a pioneering fMRI study by Kuhl et al. found reduced ACC activity over retrieval practice trials (Kuhl et al., 2007). Critically, the more that ACC activity declined, the more RIF arose for competing memories on the later retention test, suggesting that successful inhibition during retrieval practice trials reduced conflict-related activity in the ACC. Similarly, electrophysiological studies indicate that (a) increased midfrontal theta activity is associated with control demands required during retrieval of memories (Staudigl et al., 2010) and (b) decreases in mid-frontal theta power over repeated retrievals predicts greater RIF, reflecting conflict reduction benefit (Hanslmayr et al., 2010). In addition, beta band activity in the medial and lateral prefrontal regions has also been associated with inhibitory control. For example, Castiglione et al. demonstrated that inhibiting a thought from coming to mind increases beta band activity in right-dorsolateral prefrontal cortex (right-DLPFC) (Castiglione et al., 2019). Comparable results have been reproduced in the directed forgetting paradigm, a phenomenon also believed to be influenced by inhibitory mechanisms (Hubbard and Sahakyan, 2023). Beta band increases in LPFC have also been reported before successfully inhibiting a saccade (Hwang et al., 2014). A recent review paper has shed light on the domain-general aspects of inhibitory control mechanisms, suggesting that beta band activity in the frontal cortex serves as a universal marker of inhibitory control across domains (Wessel and Anderson, 2023). Together, these studies point to the involvement of medial and lateral prefrontal cortex regions in inhibitory control, with changes in theta and beta brainwave activity frequently serving as markers of this involvement.

In our study, we used anodal high-definition transcranial direct current stimulation (HD-tDCS) to investigate the causal role of the mPFC in inhibitory control during memory retrieval within a RIF paradigm. Following the study phase, we administered 2mA direct current stimulation to the mPFC for 15 minutes while participants rested. This form of stimulation is referred to as “offline stimulation”. Immediately after stimulation, participants engaged in retrieval practice tasks (repeated three times with short breaks) while their brain activity was monitored using electroencephalography (EEG). We posited that by stimulating the mPFC before engaging in retrieval practice, participants’ ability to recognize and inhibit interfering memories during the retrieval practice trials would be selectively modulated without any influence on performance in retrieval practice trials. If so, stimulation should reduce indices of RIF during the later test phase but have little effect on FAC. In addition, the disruptions in inhibitory control observed behaviourally would be accompanied by corresponding changes in the brain’s neural activity patterns, particularly in theta and beta band activity. We anticipated that these changes would specifically affect the conflict reduction benefit phenomenon, which manifests as a decline in theta band activity with repeated retrieval attempts.

## Materials and Methods

### Participants

Fifty participants (23 females, mean ± SD age, 21.4 ± 2.0 years) were recruited for the study from the student pool of the Chinese University of Hong Kong. All participants were right-handed, with normal or corrected-to-normal vision, and no reported history of psychiatric or neurological illness. Participants were briefed on the protocol, signed a written consent form, and were randomly assigned to stimulation and sham groups with no difference in age (t = 0.342, p = 0.245). All participants were paid for their participation. The study was conducted in accordance with the Helsinki declaration and was approved by the ethics committee of the Chinese University of Hong Kong.

### Retrieval Practice Paradigm

The RIF paradigm, employed in the study consisted of 24 categories, each containing 8 exemplar words in the Chinese language. Each category included four strong exemplars that were relatively easy to recall and four weak exemplars that were more challenging to remember. All exemplars were Chinese double-character words with clear meanings and were selected from word databased of Chinese speakers (Liu, 2013). In the study phase, participants studied all the items in category-exemplar word pair format (e.g. Fruit Apple). In the retrieval practice phase, participants performed retrieval practice on only the 4 weak exemplars from half (12) of the categories, with each of these items practiced three times. Retrieval practice trials presented participants with the category name and a distinctive cue for the appropriate exemplar and required participants to recall the studied item that fit those cues (e.g. Fruit Ap__). After administering a short distractor task, we then tested the category-exemplar pairs using a category-plus stem cued recall test (e.g. Fruit A__). Thus, 196 exemplars were divided into 4 types, including the practiced weak items from practiced categories (RP+ items), the non-practiced strong items from practiced categories (RP-items), the weak non-practiced items from non-practiced categories (NRP+ items), and the strong non-practiced items from non-practiced categories (NRP-items). NRP+ and NRP-trials served as control items for RP+ and RP-items, respectively.

## Experimental Design

The experiment was structured into 5 main parts, including, the study phase, electrical stimulation, retrieval practice phase, distractor phase, and test phase appearing in the order outlined in Figure 1. Further details about each of the parts are provided below.

**Figure 1.**
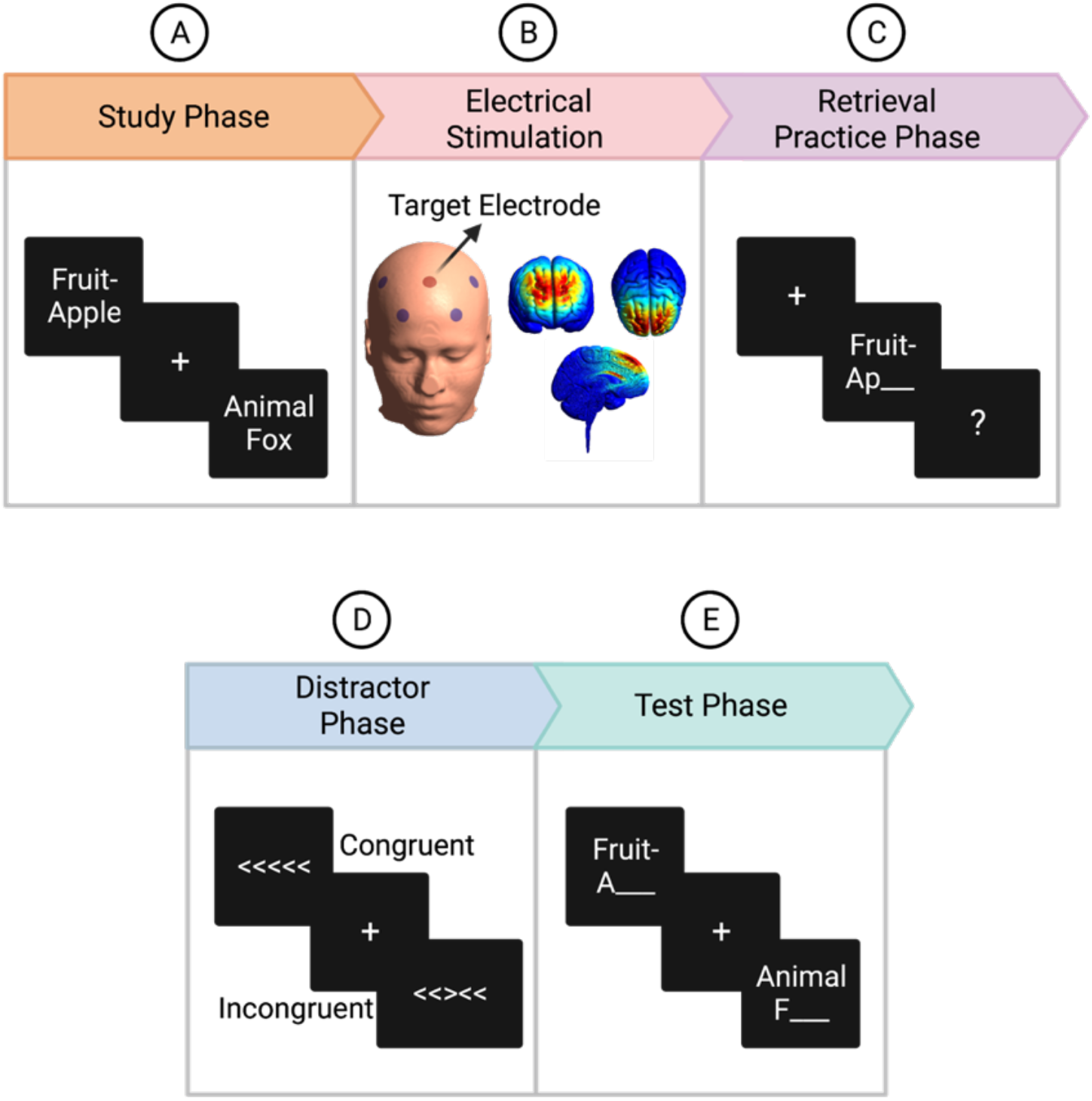
Experimental Design. **(A)** During the study phase, participants were instructed to study the items presented on the screen and were told that they would be tested later on those items. **(B)** Following the study phase, stimulation was performed for 15 minutes, during which participants rested. Stimulation electrodes were placed in a ring configuration based on the 10-20 EEG electrode placement method. The target electrode was placed at the Fz location, whereas 4 return electrodes were placed at AF3, AF4, FC3, and FC4. The simulation generated using SimNIBS is shown. **(C)**Immediately after stimulation, retrieval practice was performed in which participants recalled some of the exemplars (RP+ items) when the category cue associated with the exemplar stem appeared on the screen. Participants performed three retrieval practices for each practiced item. **(D)**Post-stimulation measurements then continued with a distractor phase lasting for 5 minutes followed by a final memory test phase **(E)** during which memory for all the studied items was tested.

### Study Phase

During the study phase, participants studied all the category-exemplar word pairs and were instructed to remember them for a later recall. We divided the pairs into 8 blocks, and each block consisted of 24 exemplars, one from each of the 24 categories. We further divided exemplars in each block into 6 sub-blocks with 4 exemplars in each (RP+, RP-, NRP+, NRP-) and exemplars within each sub-block alternated between strong and weak exemplars. We presented a filler item after each block and two buffer items at the start and the end of the experiment to control for primacy and recency effects. Trials appeared on the screen for 3 s each, separated by fixation cross for 1.5-2 s, with the whole phase lasting approximately 18 mins. Figure 1A shows the study phase presentation procedure.

### Electrical Stimulation

We stimulated mPFC with a high-definition transcranial direct current stimulator (DC-Stimulator, Sorterix Medical, Inc., NY, USA). We implanted stimulation electrodes in the EEG cap in a ring configuration. We placed the target (anode) electrode on the Fz location whereas we placed return electrodes (cathodes) at AF3, AF4, FC3, and FC4 locations based on 10-20 systems as shown in Figure 1B, thus stimulating with an excitatory protocol often termed as anodal stimulation. For the stimulation group, we administered a direct current of 2mA for 15 minutes; in contrast, for the sham group, the current ramped up to 2mA in the first 30 secs, ramped down to 0 in the next 30 secs, with no stimulation provided during the remaining 14 minutes. We applied a conductive gel to keep the impedance below 15k Ohm.

### Retrieval Practice Phase

We administered the retrieval practice phase after stimulating the mPFC for 15 minutes. We collected EEG data during this phase. Participants practiced 48 RP+ items, 4 from each of the 12 independent categories (hereinafter referred to as “practiced categories”). We divided trials into 4 randomly ordered blocks, each with 12 randomly ordered exemplars, one from each category, with a filler item appearing after each block. We repeated retrieval practice three times, each separated by a 30 s break. On each retrieval practice trial, cues appeared on the screen for 2 s, and subjects were instructed to speak the correct answer word out loud only when ‘?’ appeared on the screen. Figure 1C illustrates a typical trial sequence for the retrieval practice task. The experimenter manually recorded whether each participant’s response was ‘accurate’, ‘not accurate’, or ‘not responded”. Each of the three repetitions of the retrieval practice list lasted approximately 6 minutes.

### Distractor Phase

We conducted a 5-minute flanker task to keep participants occupied during the distractor phase (Verbruggen et al., 2006). Five arrows appeared in the centre of the screen. On congruent trials, all the arrows pointed in the same direction (left or right), whereas on incongruent trials, the central arrow pointed in the opposite direction to the surrounding flankers, as shown in Figure 1D. We asked participants to respond to the central arrow by pressing the left or right key to indicate the arrow’s direction as quickly and accurately as possible. Trials remained on the screen until the participant pressed the button. A fixation cross lasting for 1.5-2 sec appeared after each trial.

### Test Phase

We tested participants on all 196 items in the final test phase. We tested each of the 24 categories, with each category having its own block, and all RP- or NRP-items for a given category tested before all of the RP+ or NRP+ items. We ordered the categories to match the mean position of practiced and non-practiced categories in the test list. Each test trial appeared on the screen for 3 sec. We asked participants to speak the correct answer to the cues aloud whenever the cue appeared on the screen. Figure 1E illustrates a representative trial sequence for the final test task.

The experimenter manually recorded whether the response was ‘accurate’, ‘not accurate’, or ‘not responded’. The test phase lasted approximately 18 minutes.

## Behavioral Data Analysis

To test for differences in the percentage of correctly recalled items during the retrieval practice phase, we analyzed performance with a repeated-measures ANOVA using a mixed factorial design with group (stim vs. sham) as a between-subject variable and retrieval practice repetition (R1 vs. R2 vs. R3) as a within-subject variable. Given a significant main effect or interaction effect, we conducted pairwise comparisons. To test for the occurrence of RIF on the final test and its modulation by stimulation, we conducted a mixed factorial ANOVA on the percentage of items correctly recalled with group (stimulation vs. sham) as a between-subject variable and item type (RP-vs. NRP-) as a within-subject variable. We performed a similar analysis to test the impact of stimulation on FAC with group (stim vs. sham) as a between-subject variable and item type (RP+ vs. NRP+) as within-subject variables. Post-hoc comparisons were performed if any main effect or interaction arose. Finally, correlation analysis was conducted between FAC and RIF for each of the two groups using robust correlation toolbox (Pernet et al., 2012).

## Electroencephalography

### Data Acquisition

EEG was acquired using a 64-channel Neuroscan system (SynAmps2, Neuroscan Inc, Herndon, USA) at 1kHz sampling frequency with electrodes placed according to the standard 10-20 system. Two pairs of bilateral Electrooculogram (EOG) electrodes were used to collect vertical and horizontal EOG signals. A conductive gel was used to keep the impedance below 5 kOhm. An electrode between Cz and CPz was used as a reference electrode, and AFz acted as the ground electrode.

### Preprocessing

Preprocessing was performed using MATLAB R2021b (MathWorks, Inc. (2021)), EEGLab toolbox (Delorme and Makeig, 2004) and Neuroscan curry (Neuroscan, 2008). The raw data was downsampled to 250 Hz and was band-pass filtered to 1-30 Hz. Before further analysis, the stimulation electrodes were removed, and data was re-referenced using a common average reference. Vertical and horizontal EOG signals were removed from the EEG data using covariance analysis to suppress co-varying signals in each EEG channel (Semlitsch et al., 1986). The data was then segmented into 3000-msec epochs, which included a 2000 msec post-stimulus window, and an additional 1000-msec pre-stimulus window. The data was visually inspected to remove artefactual epochs.

### Time-Frequency Analysis

Time-frequency representations (TFRs) were computed using complex Morlet Wavelets ranging from 3 to 10 cycles with the Gaussian width defined as full-width at half maximum (FWHM) in the frequency domain (Cohen, 2019). TFRs were computed within the 1500 msec epoch with a pre-stimulus baseline of 500 and 1000 msec of post-stimulus period. All the trials were averaged, and baseline correction was performed using the pre-stimulus period. Cluster based permutation testing was then implemented in Fieldtrip (Oostenveld et al., 2011) to identify spatiospectral clusters showing significant group and interaction effects. For interaction effect, we subjected the difference between first and third retrieval sessions (R3 – R1) to test for group (stim vs. sham) differences, and to assess the group effect, we combined the activation data from both retrieval sessions (R1 + R3) and compared the overall activation patterns between the two groups, while controlling for multiple comparison problem. A two-sample t-test was conducted between the stimulation and sham group for each sample of the channel-time-frequency triplet.

Samples with p > 0.001 were excluded and the survived samples adjacent to each other were grouped together into clusters. The spatial constraint to include clusters in follow-up analysis was set to a minimum of two neighbouring channels. An empirical distribution of the maximum across the sum of t-values within each cluster was generated by computing the t-statistics after 1000 random Monte Carlo permutations. Observed clusters with sum of t-values having a p-value of less than 0.05 from the empirical distribution were considered significant.

### Source Analysis

The specific time–frequency windows used for subsequent source imaging were determined from the significant group and interaction effects observed after permutation analysis on the sensor level data. First, the source model was defined as a 5 mm equally spaced three dimensional grid and further warped into the standard MNI space. A single-shell head model (Nolte, 2003) was adopted based on a standard T1-weighted MRI and was co-registered to the standard electrode locations. The power in each of the significant cluster was projected onto the source space using a Dynamical Imaging of Coherent Sources (DICS) beamformer (Gross et al., 2001). DICS is a beamforming technique that uses the cross-spectral density matrix to estimate the coherence between all pairs of EEG channels. The beamformer then selects signals that are coherent at a specific frequency and multiplies them by the inverse of the lead field matrix to reconstruct the sources of the EEG activity. The power within the baseline period was also projected to the source space, and the event related desynchronization (ERD) values for specific time of interest were calculated based on the following formula:

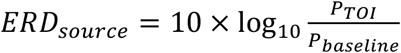

All the source analysis procedures were carried out using the FieldTrip toolbox (Oostenveld et al., 2011). Statistical analysis conducted for source-level ERD was similar to permutation. The significance level was set to p < 0.01 at the voxel level and further corrected using the Gaussian random field theory at the cluster level.

## Results

### Behavioral Results

#### No change in Retrieval Practice performance

Over the course of the three retrieval practice trials, there was a notable improvement in recall performance, indicating a significant main effect of repetition on retrieval accuracy (F (2, 96) = 64.14, p < 0.001). However, this pattern was consistent for both groups and did not exhibit an interaction between the groups (F (2, 96) = 0.96, p = 0.386), as illustrated in Figure 2A. Follow-up paired t-tests confirmed this pattern, revealing significant increases in retrieval accuracy from R1 to R2 (t (49) = 7.73, p < 0.001), from R1 to R3 (t (49) = 9.65, p < 0.001), and from R2 to R3 (t (49) = 3.80, p < 0.001). The results indicate that the stimulation of the mPFC before retrieval practice did not influence retrieval of the target memories during retrieval practice.

**Figure 2.**
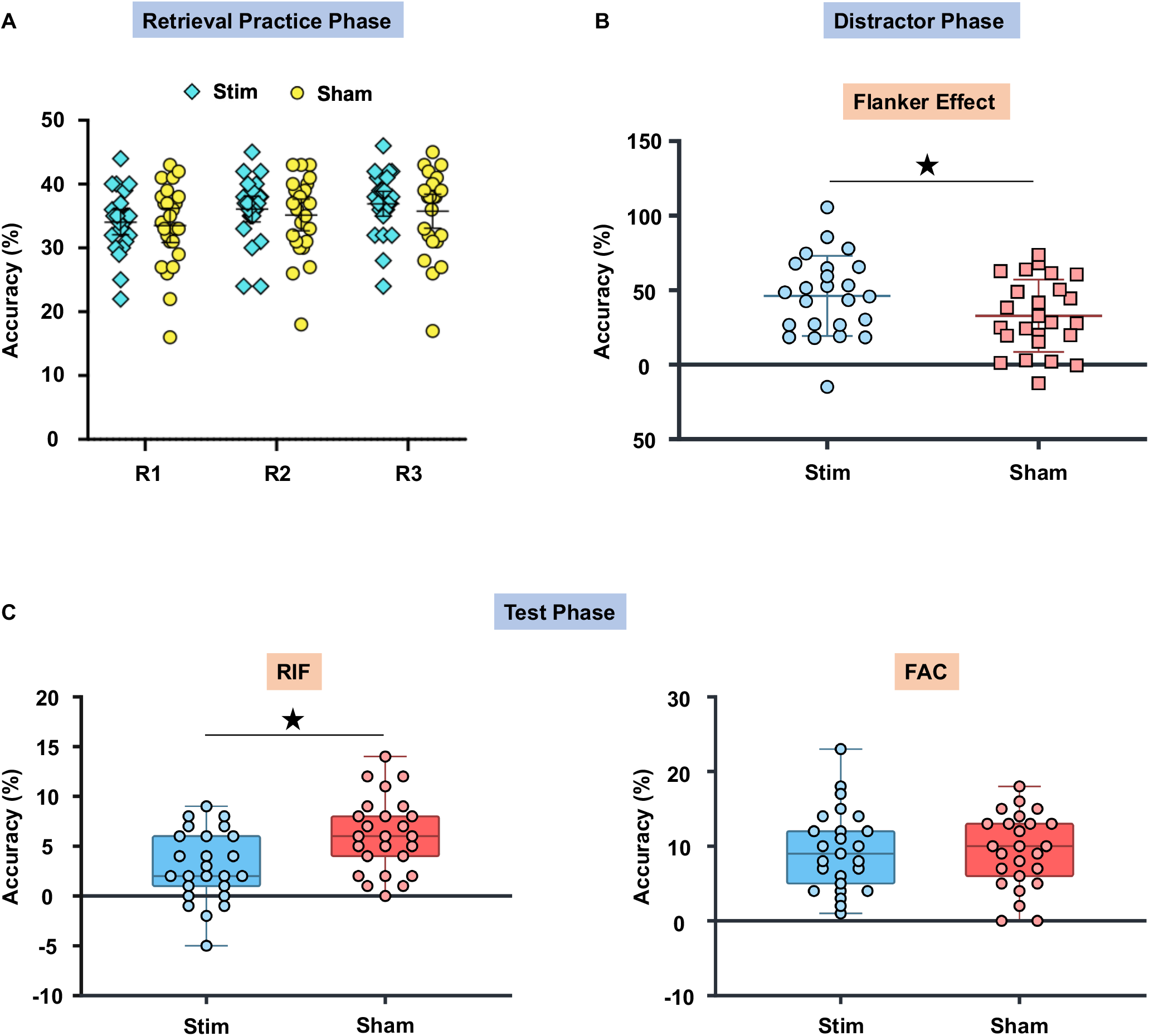
Behavioural results during retrieval, distractor, and test phase are shown. **(A)** Percentage of items correctly retrieved during repeated retrieval practice trials for the stimulation and sham groups. Accuracy for both groups increased over repetitions, and no significant interaction was observed between groups. **(B)** Stimulation impacted the performance on the Flanker task, which was administered as a distractor task immediately after retrieval practice phase. The stimulated group showed weaker ability to resist distractions compared to the sham group. **(C)** RIF and FAC effects during the final memory test phase are shown. The stimulation group showed a significantly reduced RIF compared to the sham group. FAC effect was slightly stronger in the stimulation group, but the difference was non-significant. Each dot on the box plot represents an individual participant.

#### Selective modulation of inhibitory control

Next we examined behavioural performance during the test phase. To examine stimulation induced effects on RIF, we conducted a mixed ANOVA with group (stim vs. sham) as a between-subjects factor and trial type (RP-vs. NRP-) as a within-subjects factor. The results showed a significant main effect for item type (F (1, 48) = 78.82, p < 0.001), demonstrating that the final recall of NRP-trials (Mean = 69.42 ± 8.42%) was better than the recall of RP-trials (Mean = 59.82 ± 9.55%), across both groups. More interestingly, the amount of RIF varied depending on the nature of the stimulation: we observed an interaction of item type with stimulation group (F (1, 48) = 9.07, p = 0.004) and this interaction arose due to a weaker RIF effect in the stimulation group (Mean = 6.25 ± 1.47%) than the sham group (Mean = 12.83 ± 2.03%). Notably, the RIF effect was significant in both the groups with stimulation group (t = 4.250, p < 0.001) and sham group (t = 8.215, p < 0.001) showing significant results in the t-test conducted between RP- and NRP-. The reduction in RIF in the stimulation group was driven mainly by higher recall of RP-items in that group (Mean = 62.33 ± 1.96%), compared to the sham group (Mean = 57.33 ± 1.76%), consistent with the hypothesis that stimulation disrupted the ability to inhibit those items.

In addition, this modulation was specific to RIF: FAC was uninfluenced by stimulation. A mixed ANOVA with group (stim vs sham) as between-subjects variable and trial type (RP+ vs NRP+) as a within-subjects variable showed a significant main effect for trial type (F (1, 48) = 165.84, p < 0.001). Further analyses revealed that participants recalled significantly more RP+ items (Mean = 73.08 ± 11.96%) than NRP+ items (Mean = 53.50 ± 11.00%), and this effect did not vary across the two groups (F (1, 48) = 1.32, p = 0.256). The percentages of items correctly recalled for each of the trial types are shown in Table 1 and differences in RIF and FAC are shown in Figure 2C. A correlation estimated between the amount of FAC observed for a given participant and the amount of RIF did not show significant correlations for either the stimulation (Skipped Pearson r = 0.107, CI = [-0.346 0.502], p = 0.219) or the sham groups (Skipped Pearson r = -0.129, CI = [- 0.484 0.297]).

**Table 1.** The table shows the percentage accuracy of memory recall for all the trial types (RP+, NRP+, RP-, NRP-) in the stimulation and sham groups during the test phase.

| Groups | RP+ | NRP+ | RP- | NRP- |
| --- | --- | --- | --- | --- |
| Stim | 75 ± 2.14 | 53.67 ± 2.00 | **62.33 ± 1.96** | 68.67 ± 1.96 |
| Sham | 71.17 ± 2.61 | 53.33 ± 2.42 | **57.33 ± 1.76** | 70.17 ± 1.39 |

Consistent with our hypothesis, stimulation of the mPFC impaired inhibitory control, as reflected in the reduced RIF effect in the stimulated group. Interestingly, this modulation was specific to RIF, with no change observed in FAC. This finding provides further support for the independence of RIF and FAC, as proposed by the inhibitory model, and suggests that mPFC stimulation can influence inhibitory control mechanisms.

#### Distractor Task Performance

The study employed flanker task, a classic measure of attentional control, as a distractor task administered after retrieval practice phase. The task involves participants responding to a target stimulus while ignoring conflicting cues (i.e., flankers) that surround it. A stronger flanker effect, estimated by taking the difference between reaction time of incongruent and congruent trials, indicates greater interference from the distractors, suggesting impaired inhibitory control. Our study found a marginally significant difference in flanker interference effect between the stimulation and sham groups (t = 1.83, p = 0.074). Participants in the stimulation group exhibited a more pronounced flanker effect (Mean = 46.173, SE = 5.48) compared to those in the sham group (Mean = 32.80, SE = 4.86) as shown in Figure 2B. This finding, while not the primary focus of our investigation, provides further evidence supporting the hypothesis that mPFC stimulation disrupts inhibitory control, extending beyond memory-related tasks.

### Electrophysiological Results

To examine the neural correlates of modulation of inhibitory control and identify potential differences between stimulation and sham groups, we employed a nonparametric cluster-based permutation approach on EEG data collected during the retrieval practice phase. This method is particularly well-suited for analyzing EEG data when investigating localized patterns of activation and due to its ability to handle multiple comparisons without assuming a specific distribution for the data. We were particularly interested in examining how neural activity is modulated when the retrieval cue is encountered for the first time during retrieval practice sessions and how this modulation is progressed until the final retrieval session. Thus, the analysis was restricted to only first and third retrieval sessions. For interaction effect, we subjected the difference between first and third retrieval sessions (R3 – R1) to test for group (stim vs. sham) differences, and to assess the group effect, we combined the activation data from both retrieval sessions (R1 + R3) and compared the overall activation patterns between the two groups.

#### Beta desynchronization in DLPFC indexes modulation of inhibitory control

Inhibitory effects are expected to occur shortly after the presentation of the retrieval cue, mainly driven by activity in the lateral and medial prefrontal cortices. Our group comparison of the stimulation and sham groups across the first and third retrieval sessions identified two clusters that met the criteria for significance between the groups, and both of these effects occurred within early time windows. The first cluster emerged within a brief post-stimulus time window of 0.24 to 0.32 sec in channels FC1, FCZ, C1, and CZ, coinciding with the lower beta band frequency range (15-17Hz), as depicted in Figure 3A. A broader beta desynchronization pattern can also be observed in the identified channels following stimulation in both retrieval sessions (Figure 3B) while, the cluster surviving significance threshold for group differences (stim vs. sham) is highlighted in Figure 3C. To further emphasize the observation, Figure 3D presents the TFR values extracted from the 15-17Hz range in the identified channels, indicating that beta power in the stimulation group remained low after stimulation compared to the sham group while only the highlighted region passed the significance criteria.

**Figure 3.**
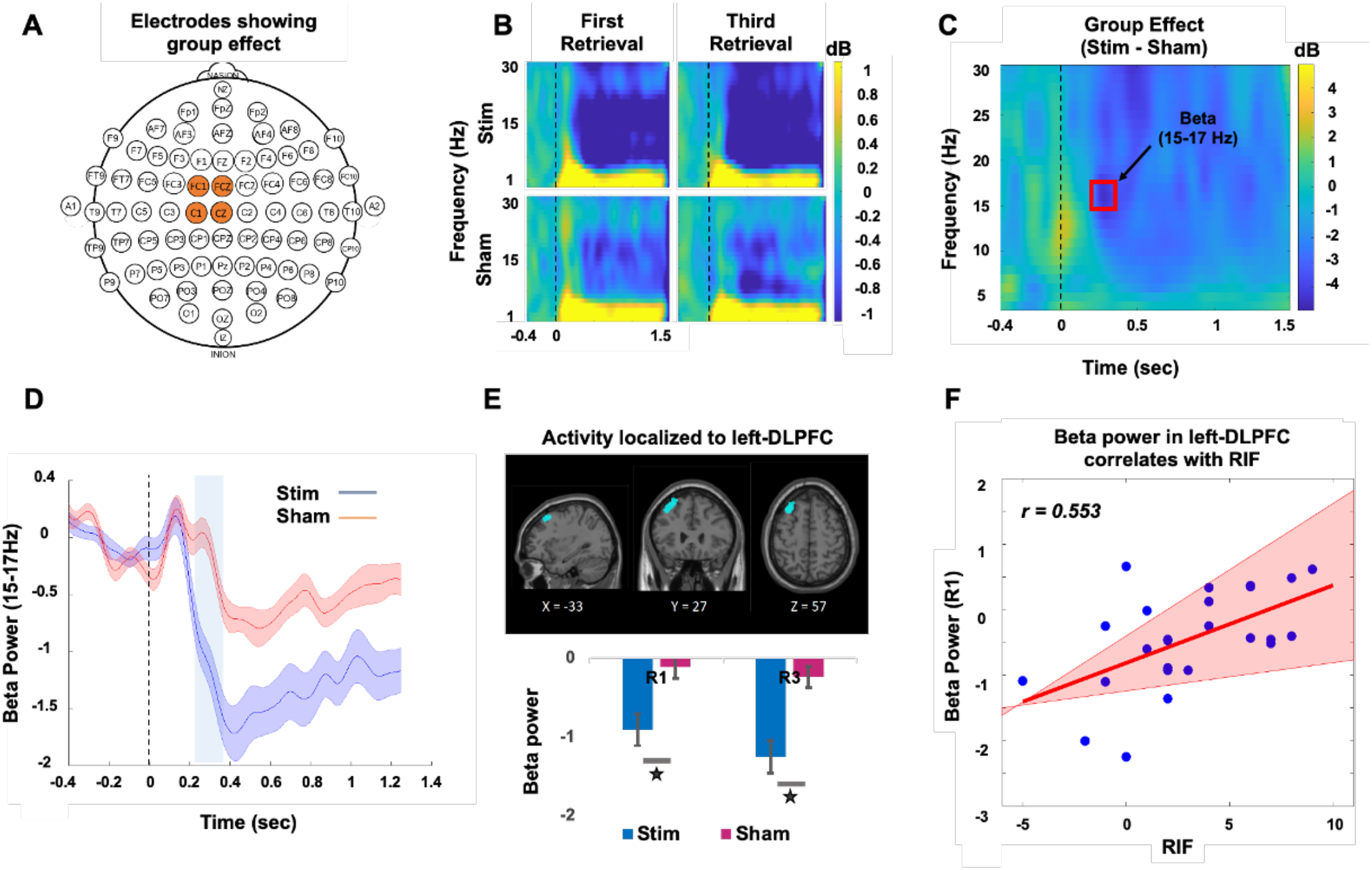
Beta desynchronization in left-DLPFC in an early time window indexes modulation of inhibitory control **(A)** Electrodes which showed a significant group effect. **(B)** Time frequency plot for first and third retrieval sessions from both stimulation and sham groups depicting a stronger beta desynchronization in the stimulated group. **(C)** The group difference between stimulation and sham group, the red box highlights the region which passed the criteria for statistical significance. **(D)** The TFR values extracted from the 15-17Hz range in the significant channels. The shaded region represents the standard error. **(E)** The observed effect in the specified time window and frequency band was localized to the left-DLPFC and the group differences were primarily driven by stronger beta desynchronization in the stimulation group compared to sham group in both retrieval sessions. **(F)** Beta desynchronization during first retrieval (R1) was directly related to RIF in the stimulation group. The red line represents the best linear fit to the non-outlying data based on Skipped Pearson correlation and the pink shaded area represents 95% bootstrapped confidence interval.

The observed group effects in the fronto-central channels support the hypothesis that the stimulation might have interfered with inhibitory mechanisms. To identify the location of this effect, we performed source localization using the TFR information i.e., time of interest (0.24 to 0.32 sec) and frequency of interest (15-17 Hz) from the first cluster. The source analysis utilized all the channels to identify significant effect in the source space using TFR information which lead to the identification of a significant effect in the left dorsolateral prefrontal cortex (left-DLPFC) as shown in Figure 3E. The beta power estimates within the left-DLPFC showed a stronger desynchronization in both retrieval sessions in the stimulation group compared to sham. We then correlated this beta-desynchronization from each of the retrieval sessions with the estimates of RIF for both the groups separately. We observed that the amount of RIF in the stimulation group strongly correlated with the beta desynchronization in the left-DLPFC during first retrieval session (Skipped Pearson r = 0.553, CI = [0.222 0.782], p = 0.004) as shown in Figure 3F, whereas no correlation was observed in the sham group (Skipped Pearson r = - 0.159, CI = [-0.580 0.381], p = 0.448). This outcome validates our hypothesis that within an early time frame, stimulation induces disturbance in inhibitory control mechanisms, with the primary involvement of the left-DLPFC in this disruption. While the correlation between RIF and beta desynchronization was evident during the first retrieval session, it was absent in the third retrieval session in both stimulation (Skipped Pearson r = -0.071, CI = [-0.334 0.453], p = 0.737) and sham groups (Skipped Pearson r = 0.205, CI =[-0.206 0.555], p = 0.325). The findings suggest that left-DLPFC driven inhibitory control mechanisms may be more specifically engaged during first retrieval compared to the third retrieval.

#### DLPFC inhibition predicts parietal beta desynchronization

The second group effect emerged again in the fronto-central channels including FC1, FCZ, FC2, C1, CZ, C2 in the upper beta frequency (23-30Hz) from 0.41 to 0.55 sec post-stimulus (see Figure 4A, 4C, and 4D). Same as cluster one there was a widespread beta desynchronization in both retrieval sessions as shown in Figure 4B. We conducted source localization on the TFRs from the second cluster and identified a source in the precuneus cortex as shown in Figure 4E. This effect did not show any correlation with RIF for either of the retrieval sessions and it possibly reflects some other neural processes related to episodic memory retrieval. Precuneus cortex has been often implicated in episodic memory retrieval processes along with contribution from inferior frontal cortex regions (Lundstrom et al., 2003; Lundstrom et al., 2005). To determine whether the early effects that we observed in beta band in the left-DLPFC relate to the later beta band effects observed in the precuneus cortex, we examined the correlation between these two regions of the brain. We found a significant correlation during the first retrieval session for the stimulation group (Pearson r = 0.462, CI = [0.067, 0.717], p = 0.020), as shown in Figure 4F. However, this correlation was not observed for the sham group (Pearson r = 0.202, CI = [-0.139, 0.509], p = 0.379). A similar but weaker correlation was also observed for the third retrieval session in the stimulation group between these regions (Pearson r = 0.360, CI = [-0.041, 0.661], p = 0.077), but not for the sham group (Pearson r = 0.112, CI = [-0.296 0.486], p = 0.559). Although our stimulation protocol was delivered to the prefrontal cortex, the observed effect was located in the precuneus. Moreover, this effect was correlated with an early left-DLPFC effect, suggesting that there is possibly network-level modulation of brain activity, through interactions between frontal and parietal brain regions. In addition, this parietal effect might relate to memory retrieval processes modulated by disruption of inhibitory control. Finally, we investigated the interaction effect by applying permutation testing on the difference of activity between first and third retrieval session. We identified a significant cluster in the parietal channels including P3, P5, PO3 in the frequency range 14-16Hz and time window of 1.256-1.328 sec (see Figure 5A, 5B, and 5C). Source localization conducted on the interaction effect TFR window localized the effect to cuneus and this interaction effect was mainly driven by a sustained beta desynchronization in the stimulation group; however, the beta power dropped during the third retrieval in the sham group as shown in Figure 5D. The beta activity for either of the retrieval sessions for each group did not correlate with RIF and neither with the beta desynchronization observed in the left-DLPFC.

**Figure 4.**
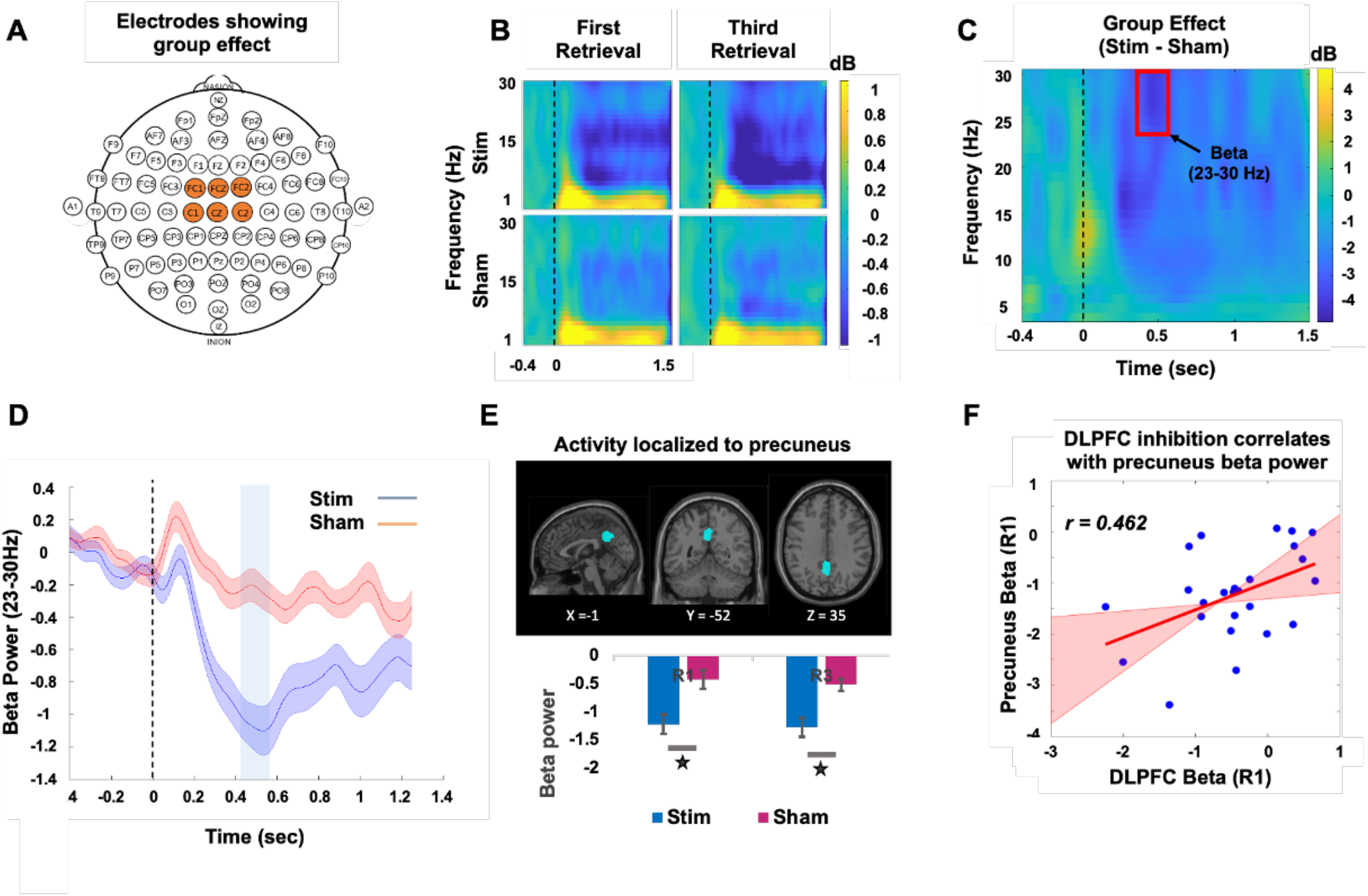
Group Effect within the frequency range of 23-30Hz and a time window spanning from 0.41 to 0.55 sec. **(A)** Electrodes demonstrating a significant group effect. **(B)** Time frequency plot for first and third retrieval sessions from both stimulation and sham groups depicting a stronger beta desynchronization in the stimulated group. **(C)** The group difference between stimulation and sham group, the red box highlights the region which passed the criteria for statistical significance. **(D)** The TFR values extracted from the 23-30 Hz range in the significant channels. The shaded region represents the standard error. **(E)** The observed effect in the specified time window and frequency band was localized to the precuneus cortex and the group differences were primarily driven by stronger beta desynchronization in the stimulation group compared to sham group in both retrieval sessions. **(F)** Beta desynchronization during first retrieval (R1) was directly related to RIF in the stimulation group. The red line represents the best linear fit to the non-outlying data based on Skipped Pearson correlation and the pink shaded area represents 95% bootstrapped confidence interval.

**Figure 5.**
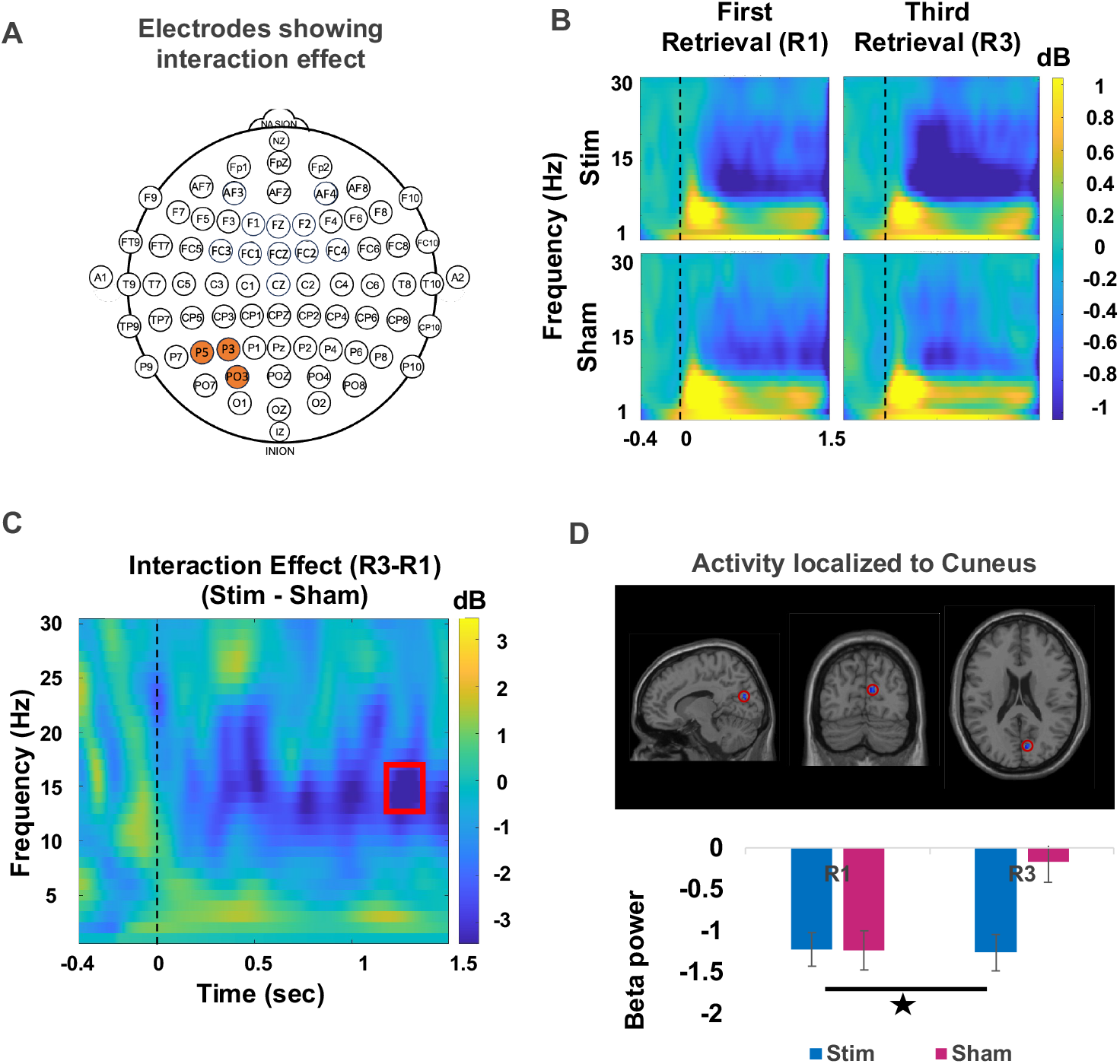
Stimulation led to an increased beta desynchronization in the parietal electrodes. **(A)** Channels which showed a significant interaction effect. **(B)** A time-frequency plot illustrating the interaction effect, with significant differences highlighted in the red box. **(C)** The source activity of the interaction effect was localized to the Cuneus. **(D)** The stimulation group showed a sustained beta power across retrieval sessions, in contrast to the declining trend observed in the sham group.

Notably, RIF did not show any correlation with any of the effects observed in the parietal cortex. This finding reinforces the hypothesis that inhibitory control is primarily mediated by the prefrontal cortex, especially the left-DLPFC, while modulation of parietal brain regions may reflect other cognitive effects related to memory retrieval, as discussed in detail in the discussion section. In addition, we would like to emphasize that there was a widespread beta desynchronization in fronto-central and parietal brain regions following stimulation. The identified group and interaction effects in very specific windows appeared only after applying strict thresholding criteria.

#### Conflict reduction benefit

One apriori hypothesis we had was that stimulation would interfere with inhibitory control by modulating theta band activity. This was based on previous findings where a decrease in theta band activity over repeated retrievals reflected the amount of RIF, often termed as conflict reduction benefit (Hanslmayr et al., 2010). Despite this, we did not observe any significant group or interaction effects in theta band activity.

We explored the possibility that the absence of modulation in the theta band could be due to the overpowering effect of beta band activity on permutation testing. To further investigate, we conducted an additional permutation test to assess group, session, and interaction effects specifically within the theta band frequency range (4-8 Hz). Our analysis did not reveal any significant clusters that met the criteria for significance for group and interaction effects (p < 0.001). However, we did observe session effects (estimated by combining exp and sham groups together and running permutation test on R1 and R3) in the theta band which showed a decrease in theta power in R3 as compared to R1. Session-effects appeared in _**R3**_ electrodes positioned over both the left and right frontal regions of the brain. The initial effect became apparent within the time ^**Sham**^ span of 0.7210 to 1.0840 sec after the onset of the event, specifically in left frontal channels F5, F7, FC5, and FT7 (Figure 6A). Subsequently, the second effect was observed between 1.0280 and 1.2480 sec, in right frontal channels F6, F8, and FC6 (Figure 6D). The time frequency plots for both first and third retrieval sessions for both the groups are shown in Figure 6B and 6E for left frontal and right frontal channels respectively. The theta power extracted from the identified left frontal and right frontal clusters is shown in Figure 6C and Figure 6F respectively, which shows a decrease in theta power in third retrieval compared to first retrieval while the highlighted region shows the area which passed the criteria for statistical significance. However, these effects did not show correlation with RIF for either of the groups.

**Figure 6.**
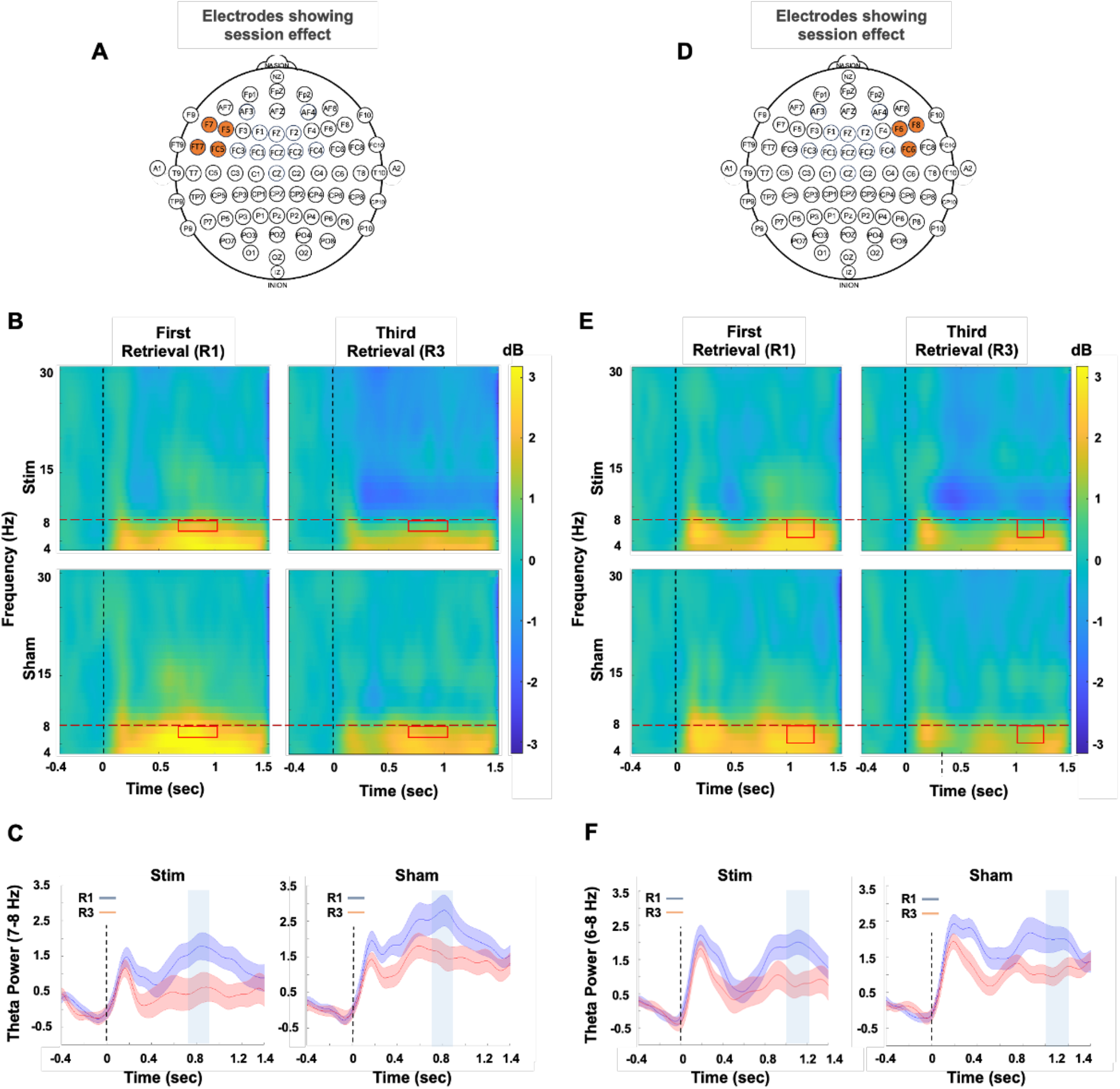
Conflict reduction benefit. Figures **(A)** and **(D)** show the channels that exhibited significant session-related effects. Figure **(B)** presents the time frequency plots of the left frontal cluster while Figure **(E)** shows it for right frontal cluster with a significant effect, highlighted in red box. The analysis was restricted to theta band (4-8 Hz), as indicated by the red horizontal dotted line. Figures **(C)** and **(F)** showcase the TFR extracted from the identified clusters within the theta frequency band in the left and right frontal electrodes, respectively. Participants in both groups demonstrated a statistically significant decline in theta power during the third retrieval compared to the first in both the identified clusters within the highlighted region.

### Stimulation related side-effects

No stimulation-related side effects were observed for any of the measured variables assessed using unpaired t-test, including headache (t (48) = 0.34, p = 0.561), neck pain (t (48) = 0.34, p = 0.561), scalp pain (t (48) = 1.75, p = 0.192), tingling (t (48) = 0.18, p = 0.677), itching (t (48) = 2.79, p = 0.102), burning sensation (t (48) = 0.28, p = 0.602), sleepiness (t (48) = 0.23, p = 0.631), concentration (t (48) = 2.24, p = 0.141) and mood (t (48) = 0.00, p = 1.000). One participant reported feeling tired after the experiment.

## Discussion

The current study employed brain stimulation to investigate how the mPFC causally influences inhibitory control while retrieving memories within a RIF paradigm. Additionally, we examined the electrophysiological factors that underlie the changes in inhibitory mechanisms induced by stimulation. Behavioural results demonstrated that stimulation prior to retrieval practice selectively modulated the amount of RIF observed during the final test phase, without affecting FAC in the stimulation group compared to the sham. Additionally, during retrieval practice, the stimulation led to a more pronounced beta desynchronization in the left-DLPFC during an early time window of 0.24 to 0.32 sec from the onset of the event and this effect showed a significant correlation with RIF. Furthermore, participants in the stimulation group exhibited a stronger beta desynchronization in the parietal cortex that sustained over repeated retrievals.

As expected, consistent with prior work, we found that retrieval practice of target memories induces forgetting of non-target competitive memories ((Anderson et al., 1994; Hanslmayr et al., 2010; Staudigl et al., 2010); see (Anderson and Hulbert, 2020; Marsh and Anderson, 2022) for reviews). Importantly, however, we demonstrate that stimulating mPFC prior to selective retrieval practice decreases RIF compared to that observed in a group receiving sham stimulation. Reduced RIF was mainly driven by higher final recall for non-target competing items (Rp-items), indicating that stimulation of mPFC reduced the tendency to forget these key items after retrieval practice. This finding is consistent with the possibility that stimulating mPFC disrupted some aspect of an inhibitory control process reliant on mPFC, preventing inhibition from suppressing competing items during retrieval practice and reducing RIF. Disrupted RIF was observed despite comparable performance in retrieving target items during the final test recall (FAC) across the stimulation and sham groups and despite improved retrieval practice accuracy over repetitions during retrieval practice phase in both groups. Supporting the inhibitory model of RIF, our data indicate that forgetting arises as a result of retrieving memories, however, these two processes are functionally independent of each other.

Other psychological manipulations have also been found to selectively disrupt RIF, without affecting FAC or retrieval practice success. For example, inducing stress before retrieval practice selectively abolished RIF with no effect on retrieval practice accuracy or FAC during the final memory recall (Koessler et al., 2009). A study by Kuhbandner et al. investigated whether affective states experienced during retrieval practice would influence RIF (Bäuml and Kuhbandner, 2007). Inducing negative moods abolished RIF, whereas inducing positive and neutral moods had no impact on RIF. It is important to note that the participants in our study did not report any mood alterations following stimulation, indicating that the observed RIF effect is not associated to mood changes. Selective disruption of RIF has also been observed after stimulation of right-DLPFC. For example, both (Penolazzi et al., 2014) and (Stramaccia et al., 2017) found that cathodal stimulation of right-DLPFC diminished RIF on a later episodic memory test, consistent with the possibility that disruption of this region compromised inhibitory control.

Whereas our study shows that stimulating mPFC with anodal stimulation disrupted inhibitory control, other studies have often reported conflicting results. For example, To et al. reported that anodal stimulation to mPFC improves inhibitory control in cognitive stroop task whereas cathodal stimulation improves inhibitory control in emotional stroop task (To et al., 2018). In another recent study, Khan et al. (2022) found that anodal HD-tDCS of the mPFC during a behavioural task improved inhibitory control (Khan et al., 2022). However, in a separate study using fMRI to investigate the impact of stimulation on the mPFC using conventional tDCS, Khan et al. (2020) observed no modulation in inhibitory control, but reported a significant reduction in connectivity between the ACC and insular cortex regions as a result of stimulation (Khan et al., 2020). To the extent that the current reduction in RIF after stimulation of mPFC reflects the disruption of inhibitory control, our findings contradict this overall pattern. One possible explanation for the discrepancy concerns the timing of the stimulation. Numerous findings suggest that the effects of online and offline stimulation can differ strikingly (Stagg and Nitsche, 2011; Pirulli et al., 2013). For example, Pirulli et al. reported contrasting effects between online and offline stimulation and observed that the effect of tDCS is more prominent when applied offline (Pirulli et al., 2013). Alternatively, the inhibitory control in attention demanding tasks, as observed in the Stroop effect, and memory inhibition, as assessed through the RIF paradigm, might engage distinct sets of cognitive processes that respond differently to direct current stimulation. Nevertheless, the inhibitory effect was similarly evident in the flanker task in our experiment, which was used as a distractor following retrieval practice. This finding casts doubt on the possibility that the inhibitory control mechanisms involved in attention-demanding tasks like the stroop or flanker tasks differ fundamentally from the inhibitory mechanisms engaged in memory retrieval.

Additionally, theta power has been traditionally associated with cognitive processing, and the midfrontal theta is recognized as an important marker for inhibitory control in attention and memory-related contexts (Hanslmayr et al., 2010; Staudigl et al., 2010; Kim et al., 2014). According to the conflict monitoring theory of attentional control, ACC detects interference and then engages DLPFC to resolve it, and this conflict detection may be partially indexed by mid-frontal theta activity (Hanslmayr et al., 2008; Kim et al., 2014; Crespo-García et al., 2022). In the context of RIF, it has been demonstrated that larger decreases in theta power over repeated retrieval practice trials predicts increased RIF observed in a subsequent test, suggesting decreased cognitive demand with repeated practice, often termed as conflict reduction benefit ((Hanslmayr et al., 2010); see (Anderson and Hulbert, 2020) for a review). Replicating prior work, here we observed a significant decrease in theta power in the third retrieval practice compared to the first retrieval in left and right prefrontal regions in both the real and sham stimulation groups. Unexpectedly, however, stimulating the mPFC with direct current did not affect overall theta power observed during retrieval practice, nor did it affect the decline in theta over practice trials. Thus, direct current stimulation before retrieval had no measurable impact on mid-frontal theta activity during the retrieval task. Given this observation, the disruption of inhibitory control observed in our study more likely derives from its impact on other neural processes not mediated by theta activity.

The other neural marker in the frontal cortex which has often been associated with inhibition is beta band activity (Hwang et al., 2014; Castiglione et al., 2019; Hubbard and Sahakyan, 2023). In our study, we observed a stronger desynchronization in the beta frequency range in the stimulation group compared to the sham group during retrieval sessions. This effect was particularly evident in the fronto-central and parietal channels, where two clusters of electrodes met the criteria for a significant group effect and one cluster for a significant interaction effect. After analyzing the group effects using source localization, two distinct sources of brain activity were identified. The first source was located in the left-DLPFC and was active during the early time window of 0.24-0.32 sec. The second source was located in the precuneus cortex and was active during the time window of 0.41-0.52 sec. Importantly, the enhanced beta desynchronization in the left-DLPFC exhibited a significant correlation with RIF in the stimulation group, indicating that participants who had a larger beta desynchronization showed lesser degree of RIF. We argue that the extent to which inhibitory control is disrupted by stimulation is reflected in the level of beta band activity desynchronization. Therefore, a greater reduction in beta band activity suggests that competing memories were not successfully inhibited, resulting in a larger reduction in RIF. In the context of inhibitory control, the timing of the occurrence of this effect is crucial. This modulation is consistent with earlier research indicating that inhibition occurs early in the memory retrieval process, with the involvement of DLPFC regions (Castiglione et al., 2019; Crespo-García et al., 2022). However, we found no correlation between beta activity and RIF in the sham group, an observation that at first seems inconsistent with the hypothesis that beta reflects inhibitory control. However, it is possible that the brain recruits additional resources to compensate for the effect of stimulation, and that reflected only in the stimulated group in the form of stronger beta desynchronization.

While increased neural synchronization has been traditionally associated with successful memory retrieval, decreased synchronization in specific frequency bands, particularly alpha and beta band within the parietal cortex, is also considered a hall mark of successful memory retrieval. (Spitzer et al., 2008; Hanslmayr et al., 2012). Surprisingly, we observed a strong beta desynchronization in the parietal cortex in the stimulation group as compared to sham and this effect was source localized to the precuneus. Considering the observations presented above, the increased beta desynchronization in the precuneus could either reflect greater retrieval of retrieval practice targets or, alternatively, increased retrieval of competitors. Given that retrieval practice performance did not vary between the stimulation and sham groups, we have little overt behavioural indication that stimulation led to more target retrieval. We do, however, have behavioural evidence that stimulation reduced RIF by selectively increasing later recall of competing items, suggesting that competitors were less likely to be inhibited by retrieval of target items. This observation suggests that failure to inhibit interfering stimuli permitted their unintended retrieval during those trials, decreasing the overall level of beta synchronization in the precuneus, relative to the sham condition. Here, it is worth highlighting that the precuneus, along with several other regions in the parietal cortex, forms a major node of the default mode network (DMN), a network associated with attentional mechanisms (Utevsky et al., 2014). Stimulation of left-DLPFC has been reported to modulate the activity in the parietal cortex possibly through network level modulation (van der Plas et al., 2021; Khan et al., 2023). Particularly the study by van der Plas et al. reported an enhancement in memory performance and an increase in beta desynchronization in the parietal brain regions after left-DLPFC stimulation with TMS during the encoding of memories (van der Plas et al., 2021). In light of these studies, we suggest that during retrieval of target memories, maintaining active representations of non-competitive memories demands increased attentional resources and this enhanced attentional engagement could manifest as increased beta desynchronization in the parietal cortex. Nevertheless, these possibilities remain speculative, and await further corroboration.

In conclusion, the study demonstrates that direct current stimulation of the mPFC selectively impedes inhibitory control processes during memory retrieval, disrupting active forgetting of competing memories. We further show that while forgetting occurs as a result of retrieving certain memories, both processes are functionally independent of each other. Moreover, stimulation also disrupted performance on a flanker interference task, suggesting that the inhibitory control process engaged during memory retrieval may be shared with the one thought to be engaged in flanker interference. Importantly, the decrease in memory inhibition was associated with a more pronounced desynchronization of beta band activity (15-17Hz) in the left-DLPFC during an early time window (0.24 to 0.32 sec) of retrieval practice. In a later time window during retrieval practice blocks, the stimulation group exhibited sustained stronger beta desynchronization in the parietal cortex, possibly reflecting retrieval of more non-competitive memories. Together, the findings demonstrate that stimulation of the mPFC with direct current disrupts inhibitory control by inducing beta band desynchronization in the fronto-parietal brain regions, and beta band desynchronization in the left-DLPFC predicts the extent of reduction in inhibition.

## Acknowledgments

We thank Golan Karvat for providing valuable comments on the draft of this manuscript. The study was supported by Hong Kong Research Grant Council (GRF No: 14205419), Hong Kong SAR, China, and UK Medical Research Council grant MC-A060-5PR00 to M.C.A.

## References

Anderson MC (2003) Rethinking interference theory: Executive control and the mechanisms of forgetting. Journal of memory and language 49:415–445.

Anderson MC, Hulbert JC (2020) Active forgetting: Adaptation of memory by prefrontal control. Annual Review of Psychology 72.

Anderson MC, Bjork RA, Bjork EL (1994) Remembering can cause forgetting: retrieval dynamics in long-term memory. Journal of Experimental Psychology: Learning, Memory, and Cognition 20:1063.

Bäuml K-H, Kuhbandner C (2007) Remembering can cause forgetting—but not in negative moods. Psychological Science 18:111–115.

Castiglione A, Wagner J, Anderson M, Aron AR (2019) Preventing a Thought from Coming to Mind Elicits Increased Right Frontal Beta Just as Stopping Action Does. Cereb Cortex 29:2160–2172.

Cohen MX (2019) A better way to define and describe Morlet wavelets for time-frequency analysis. NeuroImage 199:81–86.

Crespo-García M, Wang Y, Jiang M, Anderson MC, Lei X (2022) Anterior cingulate cortex signals the need to control intrusive thoughts during motivated forgetting. Journal of Neuroscience 42:4342–4359.

Delorme A, Makeig S (2004) EEGLAB: an open source toolbox for analysis of single-trial EEG dynamics including independent component analysis. Journal of neuroscience methods 134:9–21.

Gross J, Kujala J, Hämäläinen M, Timmermann L, Schnitzler A, Salmelin R (2001) Dynamic imaging of coherent sources: studying neural interactions in the human brain. Proceedings of the National Academy of Sciences 98:694–699.

Hanslmayr S, Staudigl T, Fellner M-C (2012) Oscillatory power decreases and long-term memory: the information via desynchronization hypothesis. Frontiers in human neuroscience 6:74.

Hanslmayr S, Staudigl T, Aslan A, Bäuml K-H (2010) Theta oscillations predict the detrimental effects of memory retrieval. Cognitive, Affective, & Behavioral Neuroscience 10:329–338.

Hanslmayr S, Pastötter B, Bäuml K-H, Gruber S, Wimber M, Klimesch W (2008) The electrophysiological dynamics of interference during the Stroop task. Journal of cognitive neuroscience 20:215–225.

Hubbard RJ, Sahakyan L (2023) Differential Recruitment of Inhibitory Control Processes by Directed Forgetting and Thought Substitution. J Neurosci 43:1963–1975.

Hwang K, Ghuman AS, Manoach DS, Jones SR, Luna B (2014) Cortical neurodynamics of inhibitory control. Journal of Neuroscience 34:9551–9561.

Khan A, Wang X, Ti CHE, Tse C-Y, Tong RK-Y (2020) Anodal Transcranial Direct Current Stimulation of Anterior Cingulate Cortex Modulates Subcortical Brain Regions Resulting in Cognitive Enhancement. Frontiers in Human Neuroscience 14:563.

Khan A, Chen C, Eden CH, Yuan K, Tse C-Y, Lou W, Tong K-Y (2022) Impact of anodal high-definition transcranial direct current stimulation of medial prefrontal cortex on stroop task performance and its electrophysiological correlates. A pilot study. Neuroscience Research.

Khan A, Mosbacher JA, Vogel SE, Binder M, Wehovz M, Moshammer A, Halverscheid S, Pustelnik K, Nitsche MA, Tong RK, Grabner RH (2023) Modulation of resting-state networks following repetitive transcranial alternating current stimulation of the dorsolateral prefrontal cortex. Brain Struct Funct.

Kim C, Johnson NF, Gold BT (2014) Conflict adaptation in prefrontal cortex: now you see it, now you don’t. Cortex 50:76–85.

Koessler S, Engler H, Riether C, Kissler J (2009) No retrieval-induced forgetting under stress. Psychological Science 20:1356–1363.

Kuhl BA, Dudukovic NM, Kahn I, Wagner AD (2007) Decreased demands on cognitive control reveal the neural processing benefits of forgetting. Nature neuroscience 10:908–914.

Levy BJ, McVeigh ND, Marful A, Anderson MC (2007) Inhibiting your native language: The role of retrieval-induced forgetting during second-language acquisition. Psychological Science 18:29–34.

Liu X (2013) The development of retrieval-induced forgetting and its mechanism. Unpublished doctorial dissertation) Tianjin Normal University.

Lundstrom BN, Ingvar M, Petersson KM (2005) The role of precuneus and left inferior frontal cortex during source memory episodic retrieval. Neuroimage 27:824–834.

Lundstrom BN, Petersson KM, Andersson J, Johansson M, Fransson P, Ingvar M (2003) Isolating the retrieval of imagined pictures during episodic memory: activation of the left precuneus and left prefrontal cortex. Neuroimage 20:1934–1943.

Marsh LC, Anderson M (2022) Inhibition as a cause of forgetting.

Murayama K, Miyatsu T, Buchli D, Storm BC (2014) Forgetting as a consequence of retrieval: a meta-analytic review of retrieval-induced forgetting. Psychological bulletin 140:1383.

Neuroscan C (2008) CURRY 7 [computer software]. North Carolina: Compumedics USA.

Nolte G (2003) The magnetic lead field theorem in the quasi-static approximation and its use for magnetoencephalography forward calculation in realistic volume conductors. Physics in Medicine & Biology 48:3637.

Oostenveld R, Fries P, Maris E, Schoffelen J-M (2011) FieldTrip: Open Source Software for Advanced Analysis of MEG, EEG, and Invasive Electrophysiological Data. Computational intelligence and neuroscience 2011:156869.

Penolazzi B, Stramaccia DF, Braga M, Mondini S, Galfano G (2014) Human memory retrieval and inhibitory control in the brain: beyond correlational evidence. Journal of Neuroscience 34:6606–6610.

Pernet CR, Wilcox R, Rousselet GA (2012) Robust correlation analyses: false positive and power validation using a new open source matlab toolbox. Front Psychol 3:606.

Pirulli C, Fertonani A, Miniussi C (2013) The role of timing in the induction of neuromodulation in perceptual learning by transcranial electric stimulation. Brain stimulation 6:683–689.

Semlitsch HV, Anderer P, Schuster P, Presslich O (1986) A solution for reliable and valid reduction of ocular artifacts, applied to the P300 ERP. Psychophysiology 23:695–703.

Shaw JS, Bjork RA, Handal A (1995) Retrievalinduced forgetting in an eyewitnessmemory paradigm. Psychonomic bulletin & review 2:249–253.

Spitzer B, Hanslmayr S, Opitz B, Mecklinger A, Bäuml K-H (2008) Oscillatory correlates of retrieval-induced forgetting in recognition memory. Journal of Cognitive Neuroscience 21:976–990.

Stagg CJ, Nitsche MA (2011) Physiological basis of transcranial direct current stimulation. The Neuroscientist 17:37–53.

Staudigl T, Hanslmayr S, Bäuml K-HT (2010) Theta oscillations reflect the dynamics of interference in episodic memory retrieval. Journal of Neuroscience 30:11356–11362.

Storm BC, Levy BJ (2012) A progress report on the inhibitory account of retrieval-induced forgetting. Memory & Cognition 40:827–843.

Stramaccia DF, Penolazzi B, Altoè G, Galfano G (2017) TDCS over the right inferior frontal gyrus disrupts control of interference in memory: A retrieval-induced forgetting study. Neurobiology of Learning and Memory 144:114–130.

To WT, Eroh J, Hart J, Vanneste S (2018) Exploring the effects of anodal and cathodal high definition transcranial direct current stimulation targeting the dorsal anterior cingulate cortex. Scientific reports 8:1–16.

Utevsky AV, Smith DV, Huettel SA (2014) Precuneus is a functional core of the default-mode network. Journal of Neuroscience 34:932–940.

van der Plas M, Braun V, Stauch BJ, Hanslmayr S (2021) Stimulation of the left dorsolateral prefrontal cortex with slow rTMS enhances verbal memory formation. PLoS biology 19:e3001363.

Verbruggen F, Notebaert W, Liefooghe B, Vandierendonck A (2006) Stimulus-and response-conflict-induced cognitive control in the flanker task. Psychonomic Bulletin & Review 13:328–333.

Wessel JR, Anderson MC (2023) Neural mechanisms of domain-general inhibitory control. Trends in Cognitive Sciences.

